# Clustering predicted structures at the scale of the known protein universe

**DOI:** 10.1101/2023.03.09.531927

**Authors:** Inigo Barrio Hernandez, Jingi Yeo, Jürgen Jänes, Tanita Wein, Mihaly Varadi, Sameer Velankar, Pedro Beltrao, Martin Steinegger

## Abstract

Proteins are key to all cellular processes and their structure is important in understanding their function and evolution. Sequence-based predictions of protein structures have increased in accuracy with over 214 million predicted structures available in the AlphaFold database (AFDB). However, studying protein structures at this scale requires highly efficient methods. Here, we developed a structural-alignment based clustering algorithm - Foldseek cluster - that can cluster hundreds of millions of structures. Using this method we have clustered all structures in AFDB, identifying 2.27M non-singleton structural clusters, of which 31% lack annotations representing likely novel structures. Clusters without annotation tend to have few representatives covering only 4% of all proteins in the AFDB. Evolutionary analysis suggests that most clusters are ancient in origin but 4% seem species specific, representing lower quality predictions or examples of de-novo gene birth. Additionally, we show how structural comparisons can be used to predict domain families and their relationships, identifying examples of remote homology. Based on these analyses we identify several examples of human immune related proteins with remote homology in prokaryotic species which illustrates the value of this resource for studying protein function and evolution across the tree of life.

**Availability:** Methods and data are available at cluster.foldseek.com

## Introduction

Proteins are the major actors in all cellular processes, from the generation of energy to the division of the cell. Knowing their structure is relevant for studying their function, their evolution and potentially for the design of drugs. While our knowledge of protein sequences has grown dramatically over the last years, reaching over hundreds of millions of sequences, the knowledge of their 3D structures has lagged behind due to the lack of highly scalable experimental methods. Improvements in methods for predicting structure from sequence^1–3^ now allow for the scalable prediction of protein structures for the known protein universe. The AlphaFold Protein Structure Database (AlphaFold DB) is a publicly available data repository of protein structures and their confidence metrics, predicted using the AlphaFold2 AI system^1,4^. The AlphaFold predicted structures have been generally assessed to be of high quality when the confidence metrics are accounted for, despite remaining inferior to experimentally determined structures^5^. AlphaFold2 and its predicted structures have now been used for diverse applications, including studies of protein pockets^6^, prediction of structures of complexes^7,8^, studies of structural similarity^9^, novel fold predictions^10^ and even improvement of genomic annotation^11^.

The large increase in available predicted protein structures has spurred the development of more efficient computational approaches, including structural data file compressions^12^, methods for pocket predictions^13,14^ and comparison of protein structures through structural alignments. For the latter, Foldseek has been developed that can increase the speed of comparisons of structures by four to five orders of magnitudes relative to previous approaches while maintaining sensitivity^15^ that makes it possible to perform structural comparisons on a large scale. Clustering proteins by their structure is a crucial tool for analyzing structural databases as it allows to group remotely related proteins. Identifying distant relationships might provide valuable insights into protein structure evolution and function. For example, protein family analysis of the initial release of about 365,000 structures^16,17^, covering the proteomes of humans and 20 model organisms, suggested that 92% of predicted domains within this set match existing domain superfamilies. However, comparing all 214 million structures against each other using current methods would take approximately 10 years on a 64-core machine. To speed up the process of clustering amino acid sequences, a linear time algorithm Linclust^18^ has been proposed to reduce the computational time significantly. However, such methods have yet to be applied to clustering by protein structural similarity.

Here, we analyze the AlphaFold Protein Structure Database that contains predicted structures for 214 million proteins across the tree of life. To be able to explore this resource, we developed a highly scalable structure-based clustering algorithm based on Linclust^18^ that structurally aligns and clusters 52 million structures in 5 days on 64 cores. We clustered the AlphaFold structural database into 2.27 million clusters with 31% of clusters - representing 4% of protein sequences - not matching previously known structural or domain family annotations. We find that 532,478 clusters have representatives present in all of the tree of life and we find several species-specific structural clusters that could contain examples of *de novo* gene birth events. Finally, we used structural comparisons to predict domain families and their relationships identifying remote homologies that expand the evolutionary coverage of previously known families.

## Results

### Structure-based clustering of the AlphaFold Protein Structure Database

The AlphaFold DB covers over 214 million predicted protein structures and has grown in several stages (**Fig. 1A**). The initial release focused on 20 key model organisms, while subsequent updates provided predictions for the Swiss-Prot dataset of the Universal Protein Resource^19^ (UniProt) and proteomes relevant to global health, taken from priority lists compiled by the World Health Organisation. The current update covers most of the TrEMBL dataset of UniProt. AlphaFold DB parses and archives these data and makes them accessible through bulk download options, programmatic access endpoints and interactive web pages. The programmatic access, in particular, facilitated the integration of AlphaFold models into other biological data repositories, such as PDBe^20^, UniProt^19^, Pfam^21^, InterPro^22^ and Ensembl^23^.

**Figure 1.**
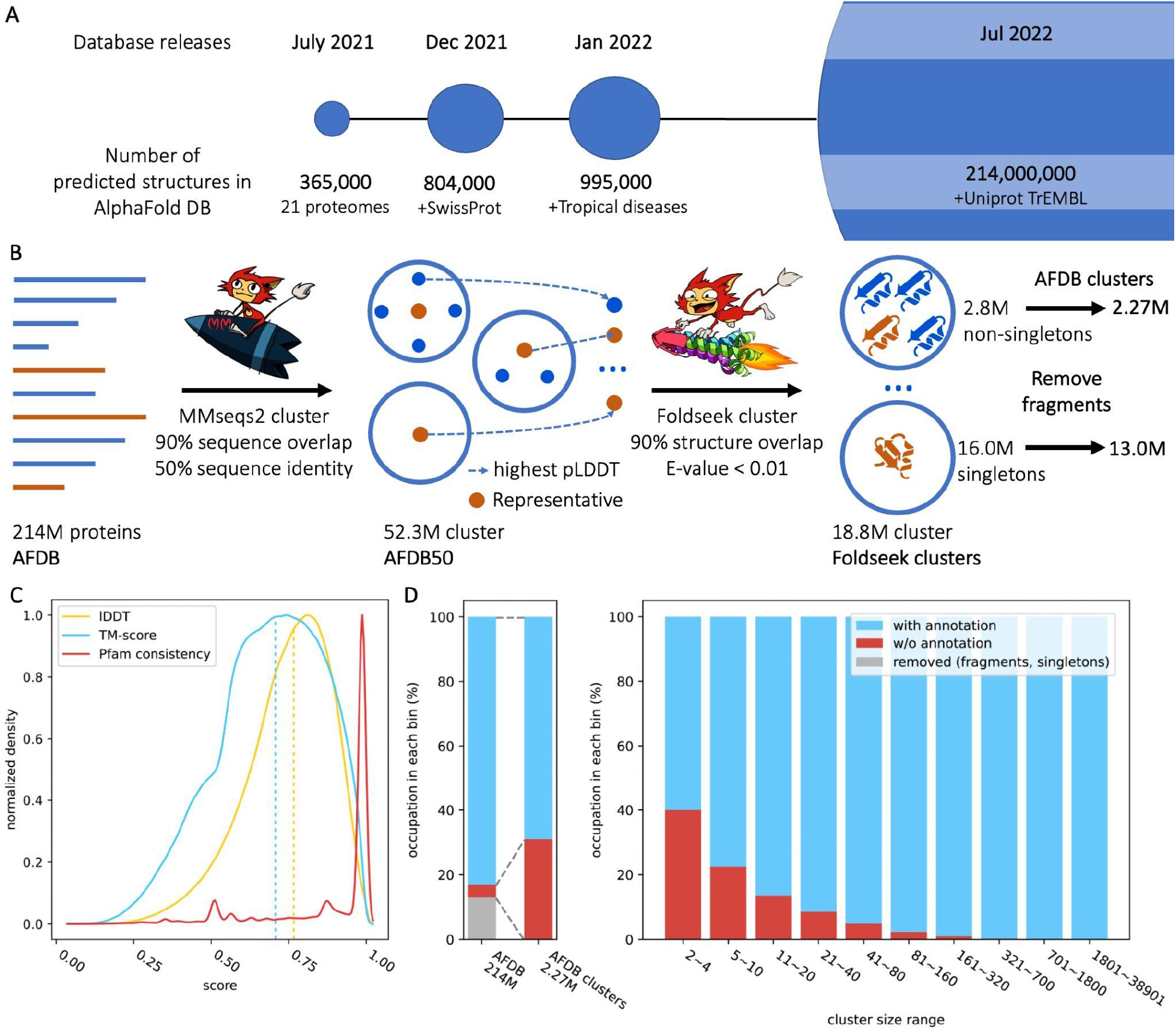
The AlphaFold database, structural clustering workflow, and summary of the clusters. **(A)** The AlphaFold DB started as a collaborative effort between EMBL-EBI and DeepMind in 2021. The database grew in multiple stages, with the latest version of 2022 containing over 214 million predicted protein structures and their confidence metrics. **(B)** A two-step approach was used to cluster proteins in the database. Firstly, MMseqs2 was utilized to cluster 214 million UniprotKB protein sequences (AFDB) based on 50% sequence identity and 90% sequence overlap, resulting in a reduction of the database size to 52 million clusters (AFDB50). For each cluster, the protein with the highest pLDDT score was selected as the representative. Next, using Foldseek, the representative structures were clustered into 18.8 million clusters (Foldseek clusters) without a sequence identity threshold, but still enforcing a 90% sequence overlap and an E-value of less than 0.01 for each structural alignment. As the last step, we remove all sequences labeled as fragments from the clustering ending up with 2.27 million clusters with at least two structures (AFDB clusters). **(C)** AFDB clusters structural and Pfam consistency. Our clusters have an average LDDT of 0.75 and an average TM-score of 0.69 across all clusters and 64% of clusters with Pfam annotations are 100% consistent. **(D)** Summary graph of sequences and clusters with and without annotation (left panel) and the relationship of cluster sizes to annotation (right panel). Each bin occupies AFDB clusters at rates of 12.0%, 10.5%, 9.2%, 10.2%, 10.5%, 10.1%, 9.1%, 9.2%, 9.2% and 9.9% from left to right respectively.

In order to gain insights into the 214,684,311 structures of the AlphaFold Uniprot v3 database we developed a scalable clustering approach in two steps as depicted in **Fig. 1B**. The first step involved using MMseqs2 ^24^ to cluster the database based on 50% sequence identity and a 90% sequence alignment overlap of both sequences, resulting in 52,327,413 clusters. For each cluster, the protein structure with the highest pLDDT score was selected as the representative. Clustering proteins by structural similarity remains computationally intensive and difficult to scale to the sizes of protein structures predicted in AlphaFold DB. For this reason, we developed a novel structure-based clustering algorithm based on Foldseek (see Methods). Briefly, we adapted Linclust and MMseqs2 sequence clustering algorithms to the 3Di structural alphabet used in Foldseek to allow structural clusters in linear time complexity. Our novel structural clustering method resulted in the identification of 18,661,407 clusters, using an Evalue of 0.01 and structural alignment overlap of 90% of both sequence criteria. It took 129 hours on 64 cores to finish the clustering. As the final step, we removed every sequence labeled as “fragment” in Uniprot. This identified 2,278,854 non-singleton clusters having on average 13.2 proteins per cluster with an average pLDDT of 71.59. The remaining 12,951,691 singleton clusters have an average pLDDT of 58.87.

We measure the quality of our AFDB clusters (**Fig. 1C**) by assessing their structural and Pfam consistency. First, we align each cluster member with a representative and calculate the average LDDT and TM-score per cluster (see Methods). The TM-score measure is based on superpositioning the structures, while LDDT is a superposition-free measure that allows for flexibility, which helps to judge the similarity of e.g. multi-domain proteins with flexible domains. Across all clusters, we found a median LDDT of 0.77 and a median TM-score of 0.71.

We also evaluate the clusters for Pfam consistency. We examine clusters with at least two sequences annotated with a Pfam domain in the UniProtKB and calculate pairwise Pfam consistency among all annotated sequences within each cluster. Our analysis revealed that 64% of these clusters have a perfect consistency score. Supplementary Figure 1 depicts the relationship between cluster members and consistency score, revealing clusters with thousands of members that have perfect consistency.

### Structurally and functionally unknown clusters in the protein universe

The availability of predicted structures covering a large fraction of the known protein universe allows us to ask what fraction of this structural space is novel. We tried to uncover structurally and functionally unknown protein clusters in the AFDB dataset - defined as “dark clusters”. We first identified 1,132,440 (50% of AFDB clusters) clusters that were found to be, at least partially, similar to previously known structures in the PDB (see Methods). Next, the representative proteins of the remaining clusters were annotated to the Pfam database by MMseqs2 search, resulting in 882,608 (39% of AFDB clusters) dark clusters (see Methods). Lastly, we identified clusters containing members with Pfam or TIGRFAM^25^ annotations in the Uniprot/TrEMBL and SwissProt database. This resulted in the identification of 705,936 (31% of AFDB clusters) dark clusters, likely enriched for novel structures.

The distribution of the known and unknown clusters as a function of their size is shown in **Fig. 1D**. The sizes of clusters that lack annotations are smaller compared to the annotated clusters. For this reason, the dark clusters map to a proportionally smaller fraction of the protein universe. While these clusters comprise approximately 31.0% of the AFDB clusters, they only represent 3.85% of the AFDB. This is inline with the expectation that structures with many representatives in the protein universe are more well studied and that the vast majority of protein structures can be annotated with at least partial similarity to a known structure of domain family annotation.

### Prediction of putative novel enzymes and small molecule binding proteins

From the 705,936 clusters without annotations (“dark clusters”), we selected 46,997 with a representative structure having an average pLDDT score of over 90 for further investigation. To focus on predicting potential novel enzymes, we searched each structure for pockets and predicted gene ontology and Enzyme Commission (EC) number using DeepFRI, a structure based function prediction method (see Methods). In total, we identified 1,723 pockets in 1,660 structures. We then predicted 5,324 functional assignments within the structures with predicted pockets. The pocket prediction led to the identification of high confidence structure predictions (pLDDT>90) that don’t appear to be correct. From 1723 pockets, 559 (32%) encompass more than 40% of the total protein (examples are shown in **SFig. 2**), indicating a general lack of compactness.

The top most often predicted molecular functions are shown in **Fig. 2A** with the top 3 including the term “transporter activity”. Similarly, the most often predicted cellular component was “intrinsic component of membrane” (391 annotations). This indicates that structures without annotations may be enriched for membrane-bound proteins which have been historically difficult to determine experimentally. Two examples of putative transporters are shown in **Fig. 2B**, including the top predicted pocket and coloured by the residue importance given by DeepFRI for this predicted function. In addition to the putative transporters, there are a wide diversity of other predicted functions. For example, A0A849ZK06 (**Fig. 2C**) is predicted as a ribonucleotide binding protein with the overall structure having an organization that resembles a protein kinase fold. The residues contributing the most to the DeepFRI prediction are directly in contact with the top scoring pocket (**Fig. 2C**), suggesting a potential nucleotide binding function for this pocket. Finally, S0EUL8 (**Fig. 2D**), has a top prediction of EC:5.6.2.-, which annotates enzymes that can alter nucleic acid conformations. The structure resembles members of the Structural Maintenance of Chromosomes (SMC) family but it is missing several characteristic elements. The preceding gene in the genome encodes for a RecN homolog (an SMC family member), giving additional evidence for a role of S0EUL8 in chromosome maintenance.

**Figure 2.**
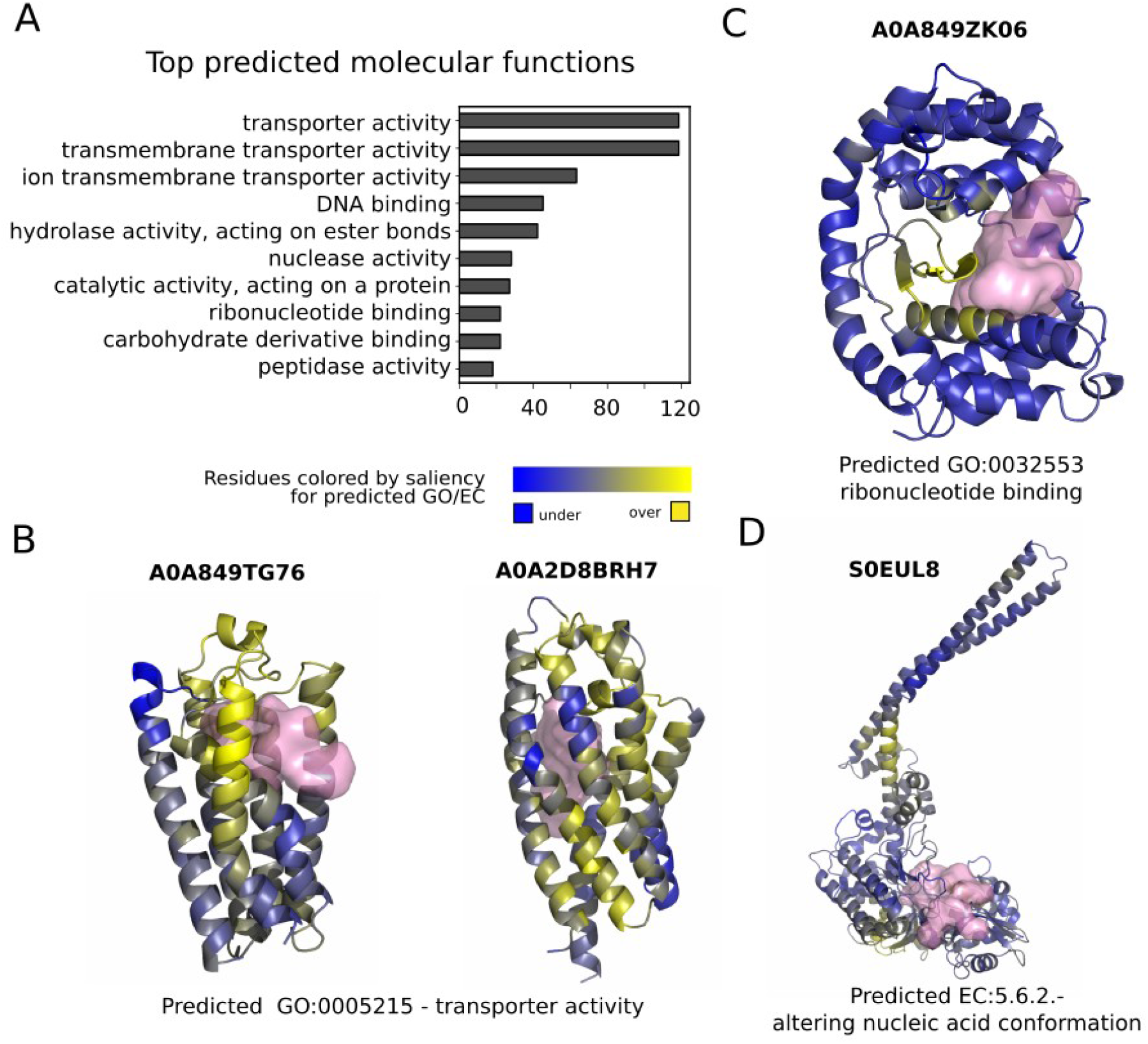
Putative novel enzymes and small molecule binding proteins in structures lacking annotation. (A) Count of Gene Ontology Molecular Function terms most often predicted by DeepFRI on the set of selected 1,660 structures with predicted pockets. (B - D) Examples of structures with predicted pockets and functional annotations. Each example shows the UniProt ID (top), highest-scoring DeepFRI function prediction (bottom) and the top-scoring pocket (pink surface). The structures are colored by residue-level contributions to the DeepFRI function predictions, ranging from blue (no contribution) to yellow (strong contribution).

### Evolutionary conservation of the structural clusters

To gain insights into the distribution of the identified structural clusters we examined their taxonomic composition to determine the extent of protein machinery shared across different super-kingdoms (see **Fig. 3A**). For this we mapped the members of the cluster in the tree of life and identified the most recent common ancestor for all members of the cluster (see Methods). Out of the non-singleton structural clusters, 532,478 (23.4%), 365,561 (16%), 307,365 (13.5%) and 11,151 (0.5%) were found to be conserved at the Cellular organism (e.g. universal to all life), Bacterial, Eukaryota, and Archaea levels, respectively. Together this suggests that the majority of the structural clusters are likely to be very ancient in origin.

**Figure 3.**
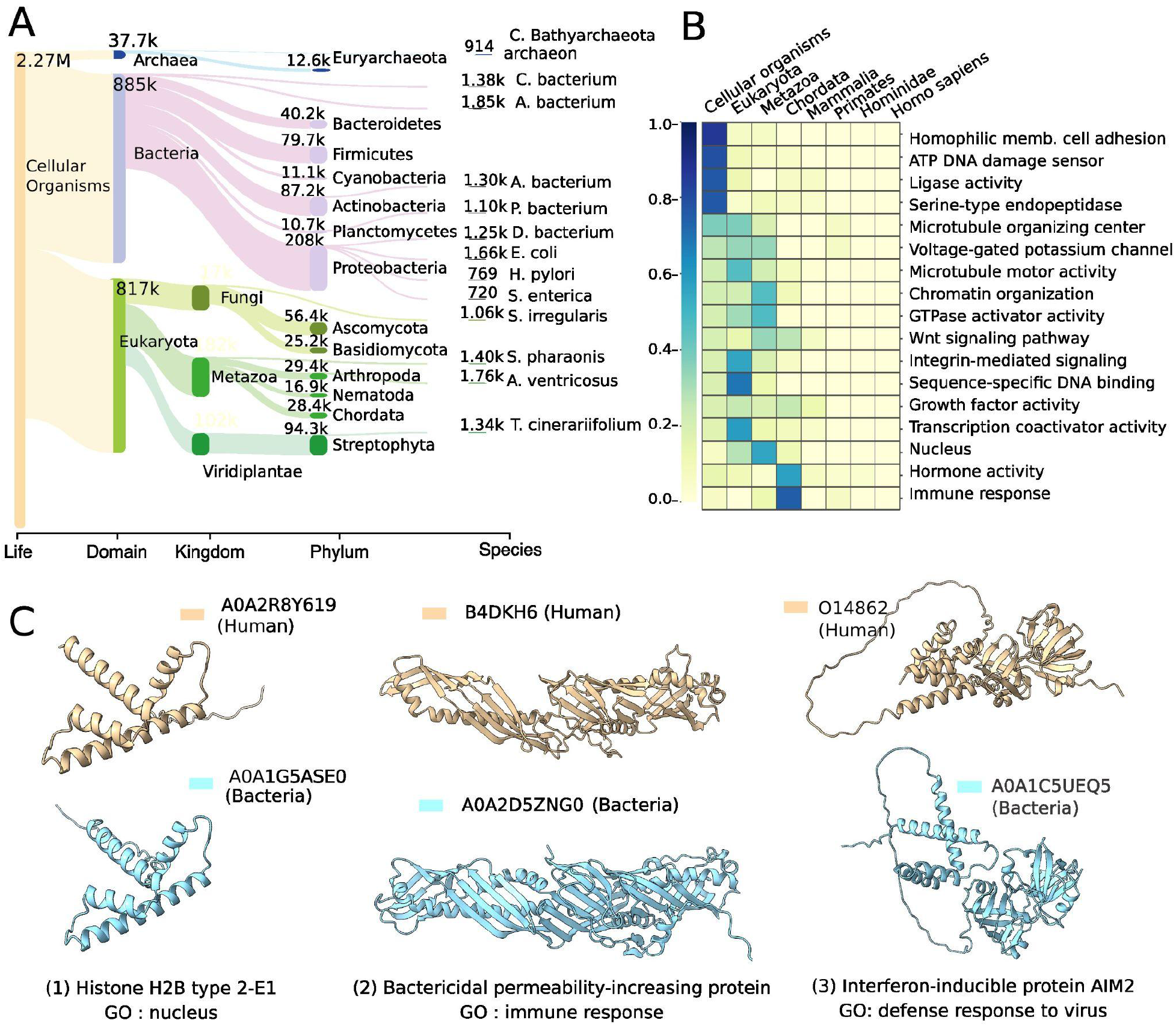
Evolutionary distribution of clusters and human-centric cluster analysis. **(A)** Visualization of the lowest common ancestor of all non-singleton clusters as a Sankey plot produced by Pavian. Only the largest 13 taxonomical nodes per rank are shown. **(B)** The distribution of selected GO terms across the human lineage is shown based on the analysis of human protein-containing clusters (abundance is normalized per GO category). **(C)** Three example structures from the human clusters that are conserved across Human and bacteria, among the eukaryote GO-annotated clusters. The left panel shows a histone protein with a nucleus GO annotation, which was found to be conserved at the cellular organism level and supports the previously reported evolutionary connection between eukaryotic and bacterial histones^26^. The middle and right panels display the human innate immunity genes BPI And AIM2 having structurally similar proteins in bacterial species highlighting the potential for cross-kingdom sharing of immunity-related proteins.

While the majority of protein clusters are mapped to the common ancestor of eukarya or older, we found 88,828 (3.8%) species-specific structural clusters. Of these species-specific clusters, 81.65% only have two members, compared to the 35.59% of clusters found to have a more ancient common ancestor. These species-specific clusters are also more likely to be “dark”, with the percentage increasing from 20.31% for those specific to cellular organisms (e.g. universal to life) to 56.62% for species-specific clusters. On average, species-specific clusters tend to be smaller proteins, with a median length of 161.5 and a mean length of 228.8, compared to the remaining clusters with a median length of 207.9 and a mean length of 281.5. Of the 43,435 members of these clusters, nearly half have an average pLDDT of less than 70, suggesting low confidence or higher disorder. However, the overall pLDDT of these clusters is comparable to that of the remaining clusters, with an average of 73.42 compared to 73.97. The organisms with the largest species-specific clusters are *Acidobacteria bacterium, Araneus ventricosus, Escherichia coli, Sepia pharaonis*, and *Chloroflexi bacterium*, which account for 1,853, 1,764, 1,646, 1,396, and 1,375 clusters, respectively.

### Evolutionary conservation of human-related structural clusters

As an example application, we studied human protein-containing clusters from an evolutionary conservation perspective. We mapped the clusters containing human proteins to the tree of life (see **SFig. 3**) and first looked for human-specific clusters (i.e. containing only human proteins). Out of the 12 human-specific clusters identified, 8 are predicted non-confident with a pLDDT score less than 70 and did not contain structural proteins. The remaining four clusters contained a herpes virus U54 (A0A126LB04) unit, Annexin (A0A4D5RA95) with limited human homologs in UniRef50, a U2 snRNP-specific A’ protein (Q9UEN1) that appeared to be a fragment but is not labeled as one, and VPS53 (A0A7P0T9Z7), a single long coil structure that was not clustered by Foldseek due to high random chances of observing such a structure. Our findings do not support the presence of newly emerging human-specific structural clusters within the set of human sequences annotated in UniProt. However, this does take into account singleton clusters.

Next, we extracted all clusters containing a human protein and associated each human cluster with its corresponding GO terms and LCA (Lowest Common Ancestor). When multiple human sequences were present in a cluster, the GO annotation of the human protein with the highest pLDDT score was selected. A small selection of GO annotations that highlight the evolutionary conservation of human structures is shown in **Fig. 3B**. Human proteins with similar structures across most of the tree of life are annotated with a diverse set of terms including several enzyme activities (e.g. ligase activity, oxidoreductase activity, serine-type endopeptidase activity). Present in bacteria and eukarya include proteins linked with the microtubule organizing center and voltage-gated potassium channel activity. Mostly restricted to eukarya include terms such as nucleus, chromatin organization and microtubule motor activity. More recently evolved structures include annotations such as immune response and hormone activity.

### Bacterial remote homology of human immunity related proteins

We noted that even if some biological processes were primarily restricted to eukarya or more recently diverged clades, we could find cluster representatives that were present in bacterial species. For example, most human proteins that are annotated to the nucleus (GO:0005634) are in clusters mapped to eukarya as their LCA. However, we find exceptions including for example a histone-related cluster (**Fig. 3C**) supporting the previously reported evolutionary link between eukaryotic and bacterial histones^26^. Similarly, we found several immunity related proteins with structural homologs present in bacteria. These include TNFRSF4 (P43489) with similar structures in bacteria due to common cysteine-rich repeat regions which overlap with the TNFR/NGFR cysteine-rich region domain annotations in InterPro (IPR001368). We also found bacterial structures related to the human CD4 like protein B4E1T0 (**SFig. 4A**) although these can also be annotated by sequence matching to the Immunoglobulin-like domain family in InterPro (IPR013783).

The structural similarity between human and bacterial proteins may also inform on their function in bacteria. The human Bactericidal permeability-increasing (BPI) protein (B4DKH6), is a key component of the innate immune system and is known to have a strong affinity for negatively charged lipopolysaccharides found in Gram-negative bacteria. In our analyses, this protein clusters with bacterial structures (**Fig 3C**), for example, the protein A0A2D5ZNG0, which aligns with the human protein at a TM-score of 0.81 normalized by the length of the human protein. Additionally, searching for partial hits by Foldseek identified the YceB from *E. coli* and other gram-negative bacteria, having structural similarity to the C-terminal region of human BPI (**SFig 4B**). The *E. coli* YceB protein is a tubular putative lipid-binding protein without a well characterized function. This structural similarity may suggest a role of YceB homologs in regulating the outer-membrane.

Our analysis identified a cluster containing the human protein, AIM2 (O14862), which recognizes pathogenic dsDNA^27^ and leads to the formation of the AIM2 inflammasome. When searching the NR database with NCBI BLAST^28^, we found no bacterial hits for the human AIM2 gene. However, three structures in Candidatus Lokiarchaeota archaeon and one in the bacterium Clostridium sp. from an uncultured source (A0A1C5UEQ5) were identified as similar to human AIM2 in our analysis. The bacterial protein (A0A1C5UEQ5), encoded on a contig of length 138,559 (GenBank FMFM01000010), is unlikely to be a contaminant due to its length^29^. A0A1C5UEQ5 is not unique, as many homologous sequences, mostly labeled as “hypothetical protein,” were found in the NR database from mostly uncultured human gut bacterial sequences with >90% sequence identity. We predicted the structure of one homologous protein that is 64% identical to A0A1C5UEQ5 (see **SFig. 5**), which originates from a cultured *Lachnospiraceae bacterium* that is part of Culturable Genome Reference^30^ of humans gut, using ColabFold^31^ and confirmed that is has a similar structure of the DNA binding domain (TMscore of 0.97 and 0.56 in relation to A0A1C5UEQ5 and human AIM respectively. These results suggest that the AIM2 inflammasome may have been repurposed from ancient DNA sensing related proteins. It is possible that the bacterial versions may also play a role in pathogen DNA sensing and response.

These results exemplify how the structural clusters can inform the evolutionary origin of specific biological processes and further illustrate the cross-kingdom similarities in immune systems.

### Prediction of domain families using structural similarity searches

The clusters defined above group structurally similar proteins at full length. Proteins are sometimes composed of different regions or domains that can fold independently, with a growing collection of such domain families being cataloged in databases such as Pfam^21^ or InterPro^22^. Domain family prediction is done primarily by sequence searches, exploring the fact that domain families have conserved sequence features. The vast increase in protein structures and fast algorithms to compare them opens the possibility of predicting domain families by structural similarity. Here, we devised a procedure using structural similarity matches by Foldseek to predict putative domain regions and families (see **Methods**, **Fig. 4A**). Briefly, a representative structure from each of the Foldseek clusters defined above was used for an all-by-all structural similarity search using Foldseek. While these representative structures should be structurally non-redundant at the full protein level, they will still share many structurally similar domains. For each sequence/structure we cluster the start and end positions of all Foldseek hits and use these to define likely domain boundaries. The predicted domain regions were then connected if they had structural similarity and a network clustering method was used to cluster domain regions into putative domain families (see Methods).

**Figure 4.**
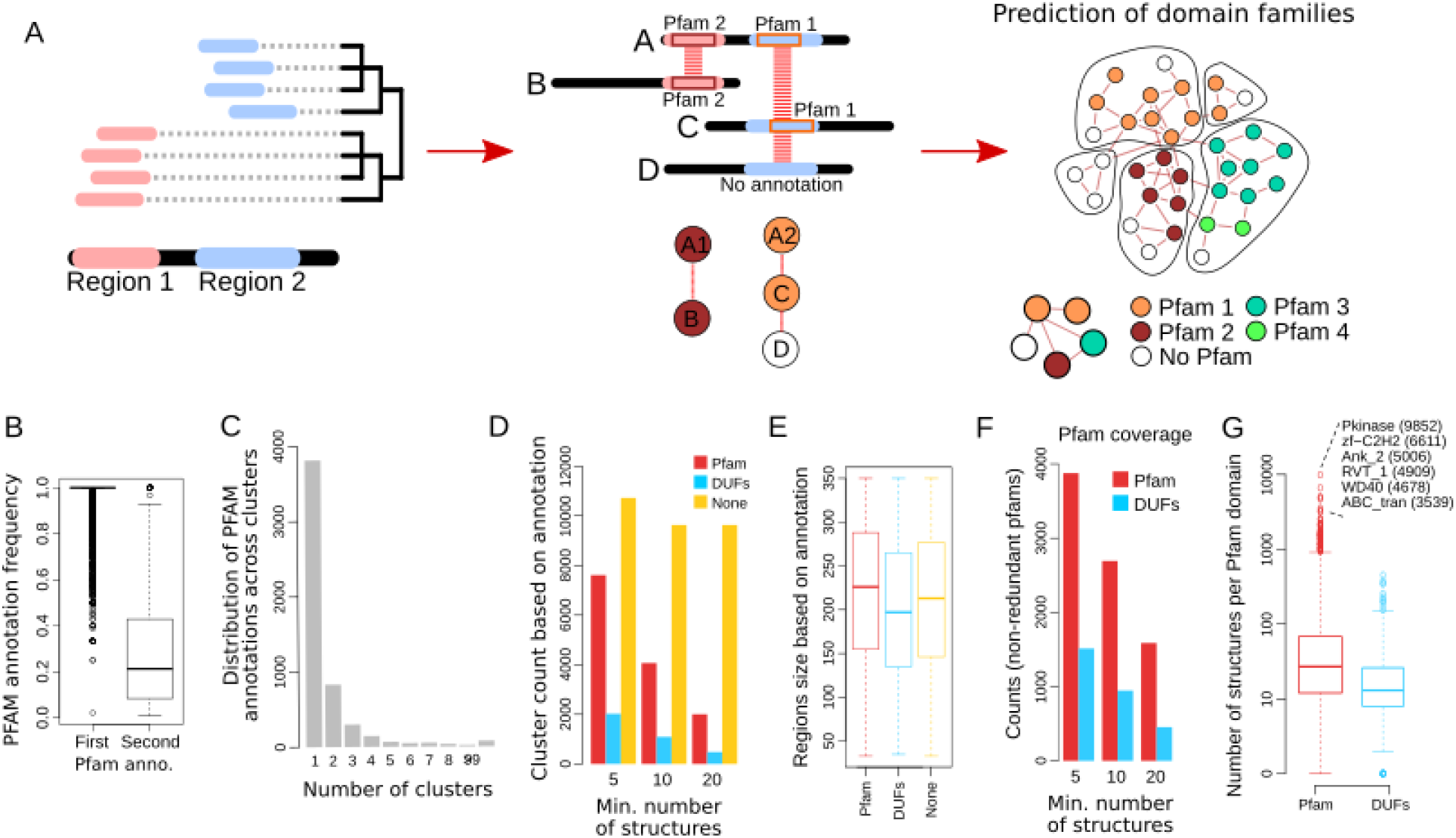
Prediction of domain families by local structural similarity hits. **(A)** Diagram illustrating the structure based domain family prediction method. Clustering of start and end positions for Foldseek hits of one protein against all others is used to define potential domain boundary positions. Each predicted domain region is linked to others sharing structural similarities and graph based clustering is used to define domain families and inter-domain similarity. **(B)** The boxplots contain the frequency distribution of the most common and second most common Pfam annotations found members of all predicted domain families. **(C)** Histogram for the counts of the number of clusters having a given Pfam as the most frequent. **(D)** Number of domain family clusters annotated to a Pfam, DUF or no domain annotation. **(E)** Distribution of protein region length in the predicted domain families, stratified by their annotations. **(F)** Non-redundant count of Pfam and DUF domain families found in the structure based predicted families. **(G)** Distribution of the number of structures found for each predicted domain family annotated with a known Pfam or DUF domain.

We used Pfam annotations to assess the quality of these predictions (**Fig. 4B-G**). For each putative domain family with ≥5 representatives we determined the frequency of the first and second most frequent Pfam annotations, with the majority having homogeneous annotations (**Fig. 4B**). Each Pfam annotation is predominantly found within a single domain family suggesting that these tend to be non-redundant. For domain families with ≥5 representatives, 7,599 families match Pfam, 2,032 match Pfam Domains of Unknown Function (DUFs) and 10,722 do not match Pfam and are likely enriched in novel families. The median length of the regions is similar for previously known or putative novel families (**Fig. 4E**). Given that we started with mostly non-redundant structures, we don’t expect this approach to recover most domain families. We found 5,388 non-redundant Pfam annotations for predicted domain families with ≥5 representatives, corresponding to ~29% of the 19,000 known Pfam families.

In summary, clustering of local Foldseek hits can accurately predict domain families leading to the prediction of many potential unexplored families. We provided a complete list of all predicted domain families in cluster.foldseek.com.

### Structural similarity across distantly related domain families

The network clustering procedure used above also allows for the identification of pairs of predicted domain families that share some structural similarity. Among such pairs, we found ~500 connections between clusters enriched with a Pfam annotation and other domains without clear annotations, providing examples of potential functional annotations. From these we focused on connected domain families enriched in proteins from different kingdoms (**Fig 5**). The Frag1 like domains exemplify the strength of structural-based similarity searching (**Fig 5A**, Frag1 like). The Frag1/DRAM/Sfk1 Pfam domain (PF10277) annotates proteins with a 6 alphahelix bundle transmembrane region that is observed in eukaryotic species. In our analysis a domain family enriched for this Pfam annotation was linked to two additional families enriched in bacterial and archeal sequences, one enriched for a domain of unknown function (DUF998/PF06197) and a second not annotated. The 3 families are structurally identical, typically forming a 6 alpha-helix bundle, despite the very low sequence similarity between the sequences forming these.

**Figure 5.**
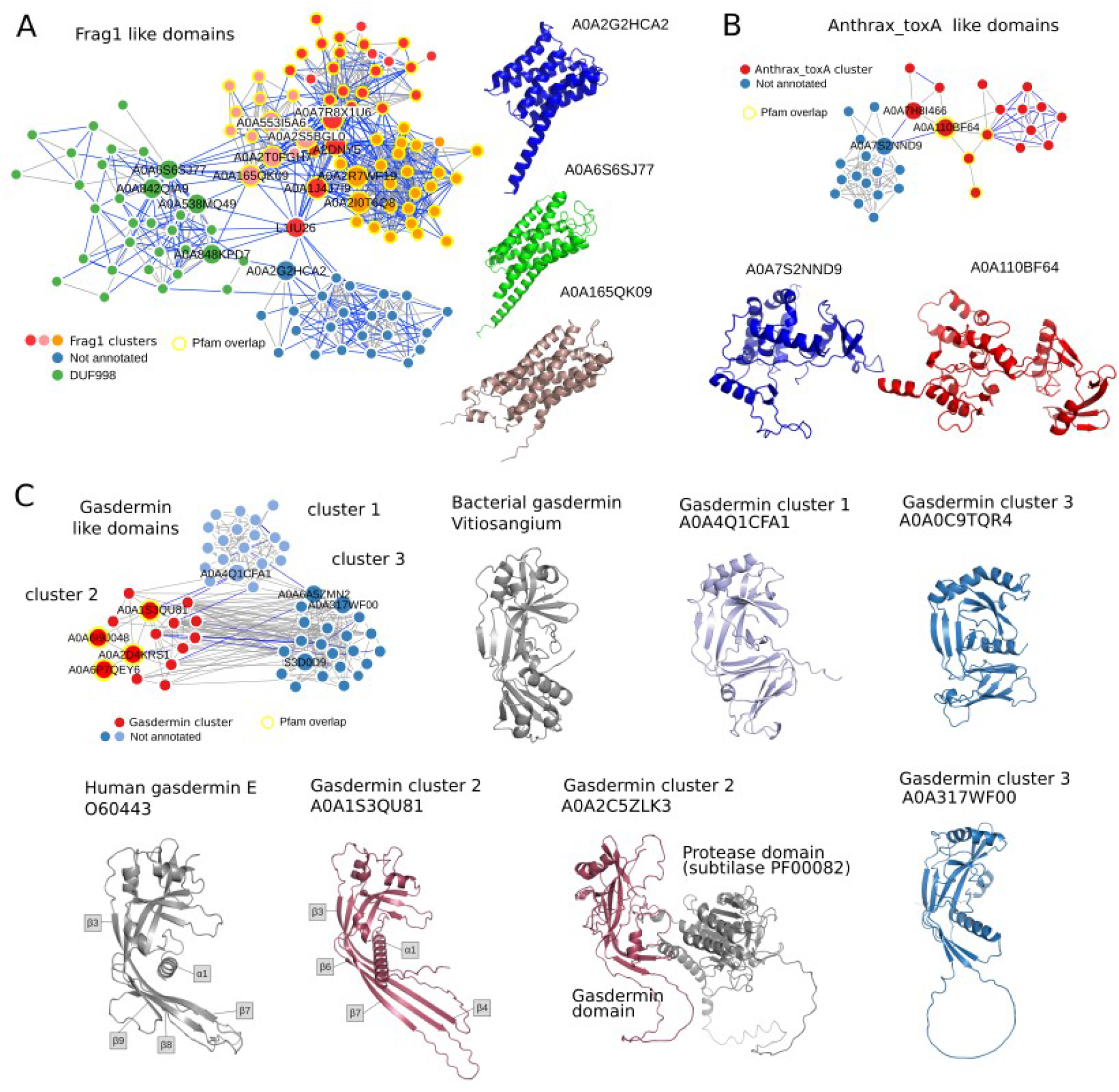
Examples of non-annotated domain families with structural similarity to annotated domain families. **(A)** Frag1 like domains, 3 clusters were found enriched for the Frag1 Pfam annotation, that had structural similarity with 1 cluster enriched for a domain of unknown significance (DUF998) and 1 cluster without annotations. **(B)** Anthrax_toxA like domains, a cluster enriched for the Anthrax_toxA Pfam annotation was found with structural similarity with a cluster having no annotations. **(C)** Two clusters without annotations were found with structural similarity with a cluster enriched for the gasdermin Pfam annotations. Human Gasdermin E and cluster 2 - Gasdermin N-terminal domain structures reveal homology to gasdermin from humans with the corresponding structural characteristics highlighted. Some gasdermin domains were found fused to protease domains (A0A2C5ZLK3). The bacterial gasdermin structure (PDB ID 7n51) is similar to novel gasdermin domains from non-annotated cluster 2. The third cluster revealed homology to both animal and bacterial gasdermins.

We also found a cluster enriched for the Anthrax_toxA Pfam (PF03497, **Fig 5B**), more specifically, the annotated domains contained structures similar to the Edema factor (EF), a calmodulin-activated adenylyl cyclase^32^. The EF is one of the 3 components forming the bacterial anthrax toxin system. Our analysis identified a structurally similar putative domain family enriched in eukaryotic proteins (**Fig 5B**). Specifically, several algae proteins were found to have structures that had partial matches to the EF domain related structures. This raises the possibility that algae might be using similar toxin systems.

### Identification of novel gasdermin like domains

Our search resulted in the discovery of 2 domain families with structural similarity with a cluster enriched for the gasdermin domain (**Fig 5C**). In human, gasdermin is the executor of inflammatory cell death called pyroptosis and is crucial for defense against pathogens. Upon sensing a pathogen, caspases are activated that cleave off the C-terminal repressor domain of gasdermin, releasing the N-terminal domain to assemble into large pores in the cell membrane^33^. The predicted gasdermin structures from all three groups exhibited the structural characteristic conservation of a twisted central antiparallel β-sheet and the shared placement of connecting helices and strands of gasdermin. The structures enriched in the gasdermin Pfam annotation adopted a similar conformation as that of the mammalian gasdermin N-terminus, especially of gasdermin E, which is considered evolutionary ancient^34^. In the inactive structure of mammalian gasdermin (A,B,D,E), the N-terminus forms interfaces with the repressor C-terminal domain mediating auto-inhibition, one of this is the primary interface at the α1 helix^35^. Gasdermin is activated by proteolytic cleavage, which results in the N-terminal activation through the lengthening of strands β3, β5, β7, and β8 and oligomerization^36^. Indeed, gasdermin domains from the Pfam annotated group had both the α1 helix as well as the corresponding β-sheets necessary for the active form of gasdermin. Gasdermin was also recently discovered in bacteria and archaea, where it is similarly activated by dedicated proteases and defends against phages by pore-mediated cell death^37^. Interestingly, the non-annotated group 1 of gasdermin domains displayed strong similarity with the bacterial gasdermin structure (**Fig 5C**). The other non-annotated group (cluster 3) showed a large degree of diversity and exhibited features of both mammalian and bacterial gasdermin. In some cases, we observed that the N-terminal gasdermin domain was fused to other domains including proteases (**Fig 5C**, A0A2C5ZLK3). As gasdermin is activated by proteolytic cleavage such protein fusion hints at a similar activation mechanism for the novel gasdermin domains.

## Discussion

The orders-of-magnitude increase in available structural models raises challenges in data management and analysis of such large volumes. For this reason, we developed a clustering procedure that can scale to hundreds of millions of structures, identifying 2.27 million nonsingleton clusters with 31% not having similarity to previously known structures or domain annotations. These clusters only annotate 4% of protein sequences indicating that the vast majority of the protein structural space has been, at least partially, annotated. As the criteria used include partial hits to known structures or domain annotations, the degree of understudied structural space is likely underestimated. As we illustrate, our analysis can guide the prioritization of predicted novel protein families for future computational and experimental characterization.

Structural clustering is a powerful tool for identifying homologous proteins, but its accuracy can be affected by certain limitations. In this study, we set a 90% alignment overlap as the requirement for assigning a structure to a cluster, which may exclude homologs with significant insertions or unique repeat arrangements. Additionally, our strict e-value threshold of 0.01 may result in missed homologs. Another limitation is that the current AlphaFold database does not contain the full extent of protein sequences from metagenomics studies or viral proteins, limiting the potential to detect retroviral proteins.

In addition to the full-length protein clustering we used Foldseek’s local hit matches to predict and cluster protein regions into putative domain families. The protein region clusters tend to overlap well with previous definitions of domain families as annotated in the Pfam database and led to the identification of over 10,000 unassigned clusters that should be enriched in putative novel domain families. We did not perform exhaustive searches with other sequence based domain family annotations that could annotate additional clusters with prior knowledge. It is important to note that we only considered the representatives of Foldseek clusters when performing the domain prediction. As the domain prediction requires multiple observations on the same structural region, additional domains are expected to be detected if each structure was searched against a larger set of structures.

As protein structure is conserved for longer periods of evolutionary time than protein sequences, we expect that AlphaFoldDB will empower the identification of remote homology. While some advanced sequence-based methods can already assist in this task^38–40^, the availability of predicted structures helps identify meaningful evolutionary relationships. Our analysis here provides several examples of structural similarity across kingdoms that is indicative of remote homology. In particular, we focused on several examples relating human immunity with bacterial structures, emphasizing how some ancient systems have been co-opted for use in the mammalian immune response system. We expect that many more examples can be derived from the clustering results provided here.

## Methods

### Structural clustering algorithm

The clustering procedure is similar to MMseqs2’s clustering but instead of using sequences, Foldseek’s 3Di alphabet (see **Fig 6**) was employed to represent the structures as 1-dimensional sequences. The clustering algorithm combines Linclust^18^ and cascaded MMseqs^41^ clustering. The pipeline applies this strategy to allow for efficient clustering of millions of structures. First, protein structures are converted to 3Di sequences and processed according to the Linclust workflow. This includes extracting m k-mers (default m=300) of length 10 from each sequence and grouping them based on their hash value. The k-mer groups are then used to assign each structure to the longest sequence (representative) within the group. The shared diagonal i-j on which the k-mer is found is also stored for further use in the alignment step.

**Figure 6.**
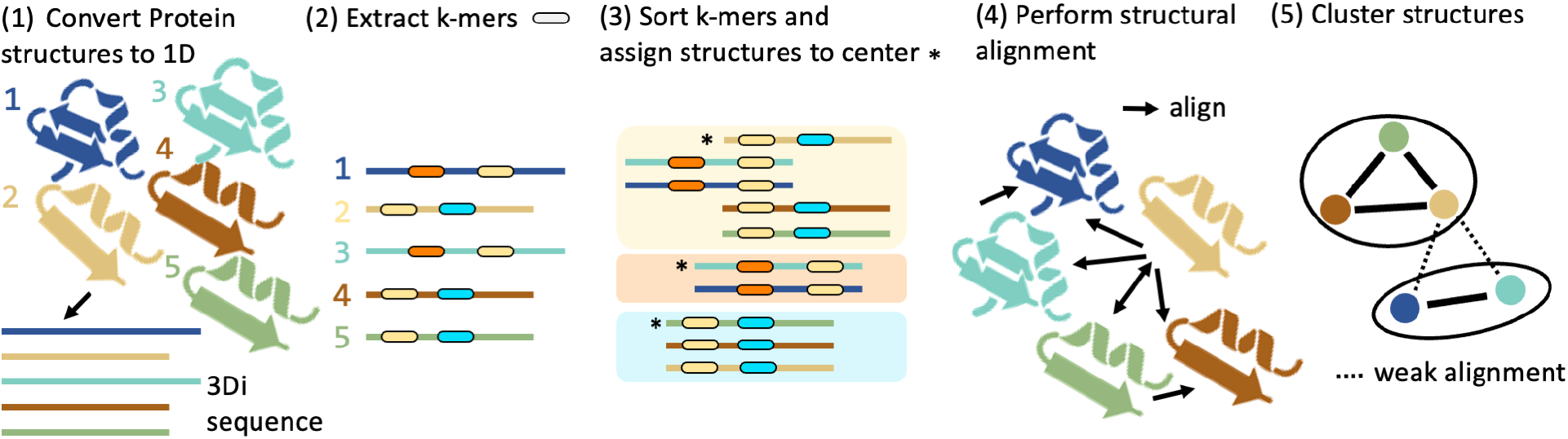
The five-step clustering pipeline for efficient clustering of millions of protein structures using Foldseek’s 3Di alphabet. (1) Protein structures are converted to 3Di sequences and processed through the Linclust workflow. (2) For each sequence 300 min-hasing k-mers are extracted and sorted. (3) The longest structure is assigned to be the center of each k-mer cluster. (4) Structural alignment is performed in two stages: first an ungapped alignment based on shared diagonal information is performed, hits are pre-clustered and second the remaining sequences are aligned using Foldseek’s structural smithwaterman. (5) The remaining structures meeting alignment criteria are clustered using MMseqs2’s clustering module. After the Linclust step the centriods are successively clustered by three cascaded steps of prefiltering, structural Smith-Waterman alignment and clustering using Foldseek’s search.

The pipeline then proceeds with an ungapped alignment algorithm that rescores the structures based on the shared diagonal between members and representatives using 3Di and amino acid information. The sequences that meet the defined alignment criteria, such as E-value, sequence identity, alignment LDDT or TMscore, are clustered using the MMseqs2 clustering module (default using the set-cover algorithm). After this step, the already assigned structures are removed from the set and the remaining representative member hits are aligned using Foldseek’s structural Gotoh-Smith-Waterman^15^, and all passing hits are clustered as well. The remaining cluster representatives are successively clustered by three cascaded steps of prefiltering, structural Smith-Waterman alignment and clustering.

### Cluster purity analysis

To assess cluster purity, we followed a two-step approach. First, we calculated the average LDDT and average TMscore per cluster to assess the structural similarity. For this, we aligned the representative to the cluster members using the “structurealign -e INF -a” module in Foldseek and reported the alignment LDDT and TMscore using --format-output lddt,alntmscore. For each cluster we compute the mean illustrated in Figure 1C.

Secondly, we evaluated the Pfam consistency of each cluster by using Pfam labels obtained from UniProt/KB. We have only taken into account the clusters that have at least two sequences with Pfam annotations and we calculated the fraction of correctly covered Pfam domains for all Pfam sequence pairs ignoring self-comparison. We define true positives as a pair of Pfam domains belonging to the same clan. For each pair, we compute the consistency scores by true positive count divided by the count of Pfams in the reference sequence. Finally, we computed the mean overall pair scores. This approach enabled us to determine the proportion of sequences within a given cluster that shared the same Pfam annotation.

### Identification of “dark” clusters and the lowest common ancestor

To eliminate clusters similar to previously known experimental structures, we conducted a search using Foldseek against the PDB (version 2022-10-14) for each cluster representative, with an e-value threshold of 0.1. We then excluded clusters annotated with Pfam domains by searching the cluster representatives using MMseqs2 with parameters -s 7.5 --max-seqs 100000 -e 0.001 against the Pfam database. Finally, we removed clusters with members annotated with PFAM or TIGRFAM20 annotations in the UniProt/TrEMBL and SwissProt database. To determine the lowest common ancestor of each cluster, we used the lca module in MMseqs2^42^ ignoring the two taxa (1) 12908 “unclassified sequences” and (2) 28384 “other sequences”. We visualized the lowest common ancestor (LCA) results using a Sankey plot generated by Pavian^43^.

### Pocket and functional activity predictions for dark clusters

We predicted small molecule binding sites for representative dark cluster members by adapting the approach from Akdel, M. *et al*. ^9^. We used AutoSite to predict pockets^44^, and selected pockets with an AutoSite empirical composite score >60 and mean pocket residue pLDDT >90 for additional analyses. To assign putative function and predict catalytic residues, we used DeepFRI^45^ to predict enriched GO/EC terms and residue-level saliency weights across available GO/EC categories (BP, CC, EC, MF). Pocket and functional predictions were then visually examined using a web app we developed (https://github.com/jurgjn/af-protein-universe).

### Domain prediction from Foldseek local hits

First, we filtered out low scoring Foldseek hits using an e-value of 10^-3^ as threshold. We defined potential domain boundary positions for each protein sequence by clustering start-stop positions (hierarchical clustering, height parameter of 250 to establish clusters). Predicted domains are then linked to others based on structural similarities, keeping the highest scores when duplicates are found. Then the resulting network is trimmed excluding connections with e-value higher than 10^-5^, predicted domains with more than 350 amino acids and connected components with less than 5 nodes. We applied graph based clustering (walktrap, 6 steps), keeping communities with at least 5 members. Each predicted domain inside the selected communities was annotated using Pfam-A regions mapped to UniProt identifiers (v35.0), more than 75% of the Pfam domain has to overlap with the predicted domain. We calculated inside each community the frequency of Pfam annotations and defined them based on the highest one. Due to its size, we decided to keep out of the following analysis one community with 152,959 structures (group ID 1;1, see supplementary files in cluster.foldseek.com). We connected the remaining communities based on the structure similarities, allowing connections with a p-value smaller than 10^-3^.

## Acknowledgments

MS acknowledges the support by the National Research Foundation of Korea, grants [2020M3-A9G7-103933, 2021-R1C1-C102065, 2021-M3A9-I4021220], Samsung DS research fund and the Creative-Pioneering Researchers Program through Seoul National University. PB is supported by the Helmut Horten Stiftung and the ETH Zurich Foundation.

## Availability

Data is freely and publicly available (CC-BY) at cluster.foldseek.com and the cluster method is available as free and open source software (GPLv3) at foldseek.com.

## Supplementary figures

**Supplementary Figure 1.**
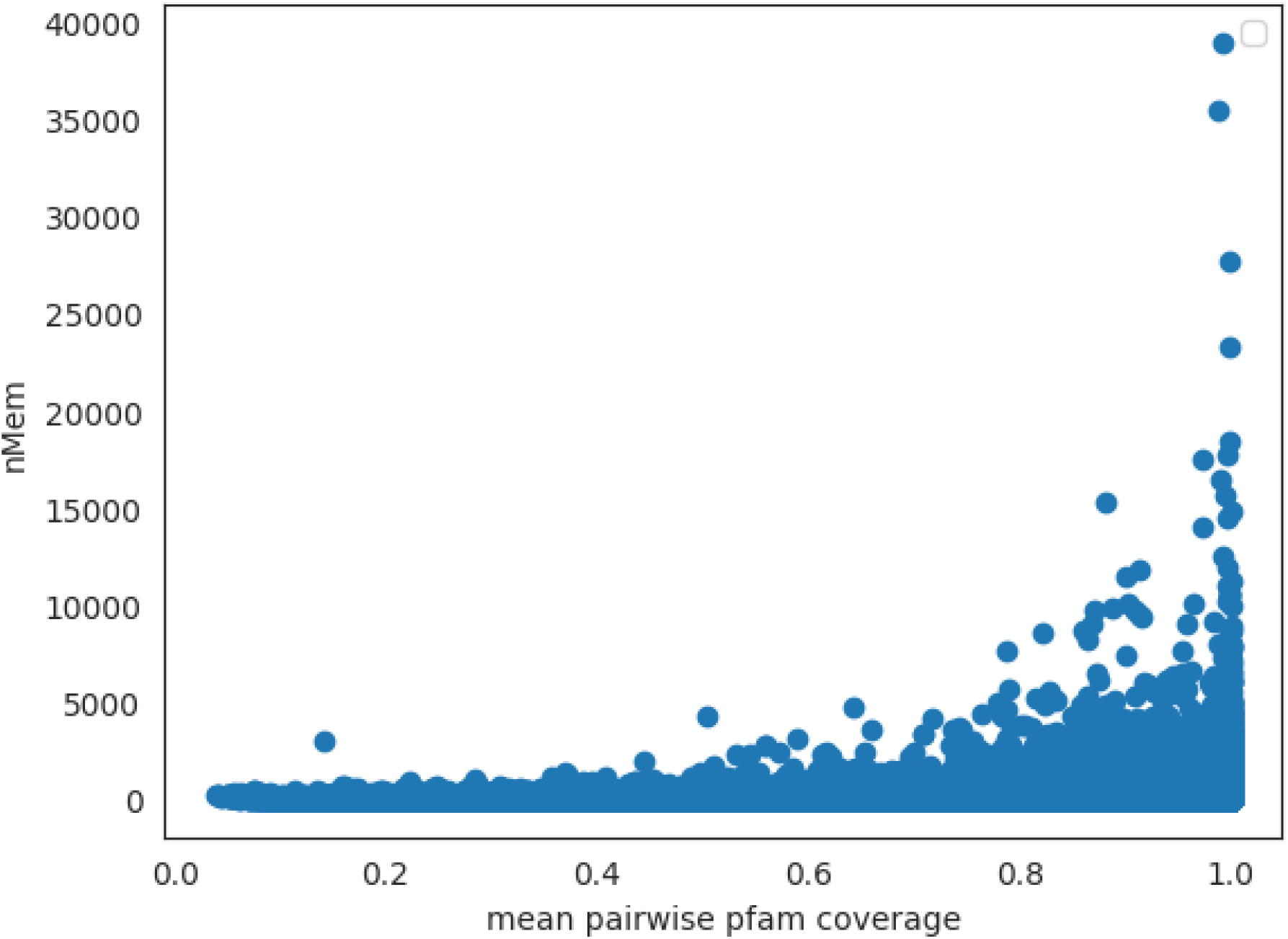
Relationship of cluster member size to mean pairwise Pfam coverage.

**Supplementary Figure 2.**
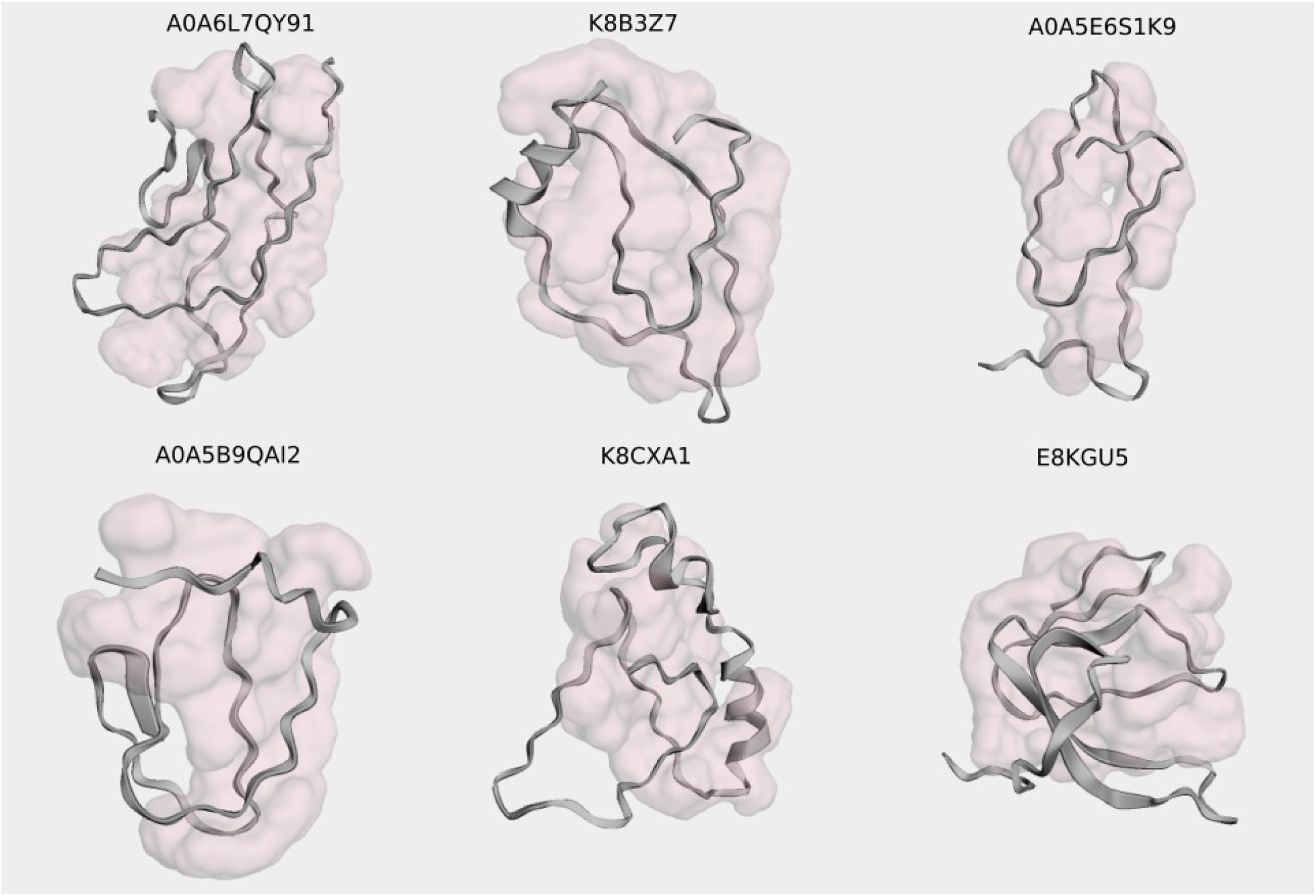
Examples of non-compact AlphaFold2 predicted structures. Examples of representative structures of clusters without annotations having pLDDT>90 and a predicted pocket covering over 80% of the residues of the structure.

**Supplementary Figure 3.**
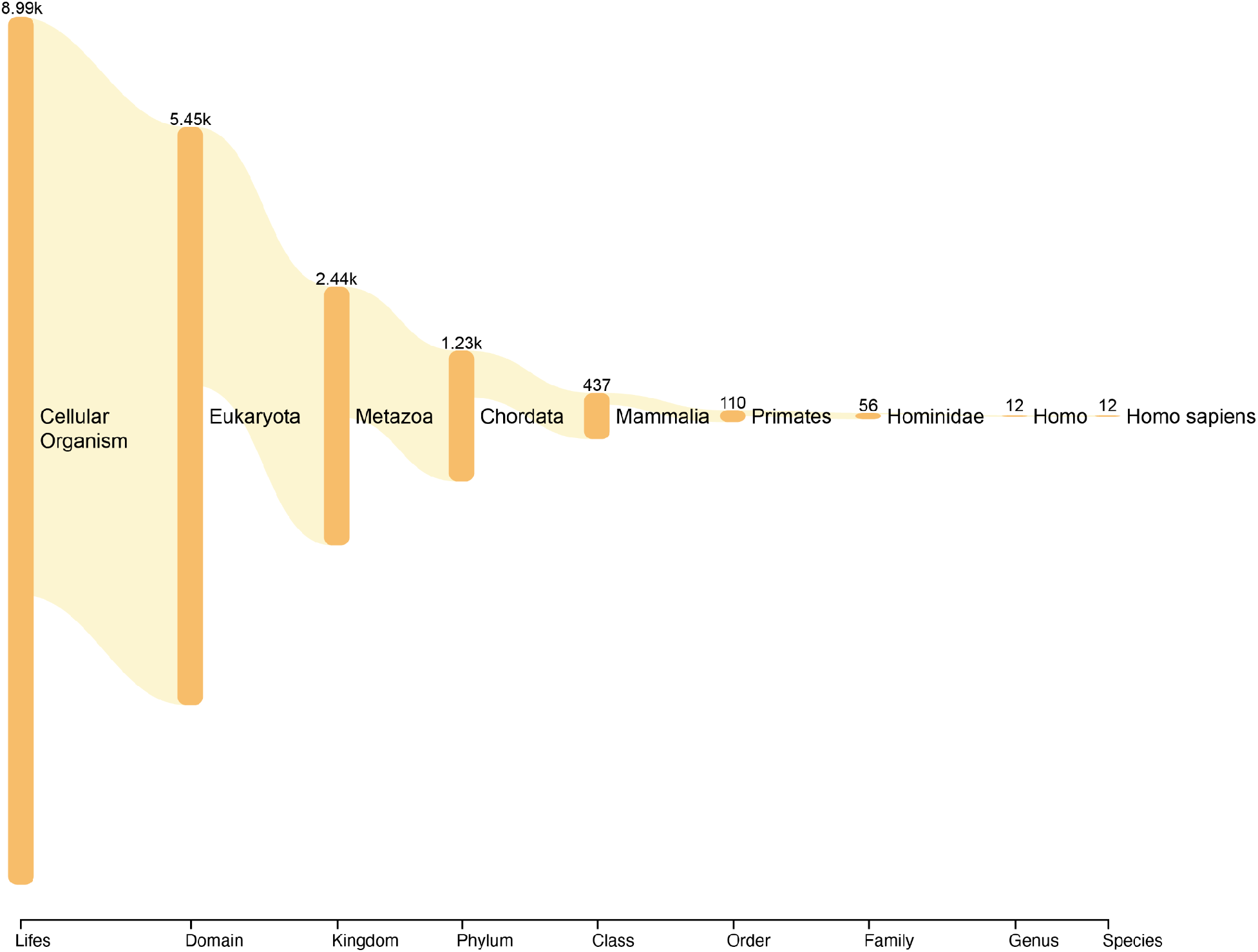
LCA plot of the clusters that contain Homo Sapiens proteins.

**Supplementary Figure 4.**
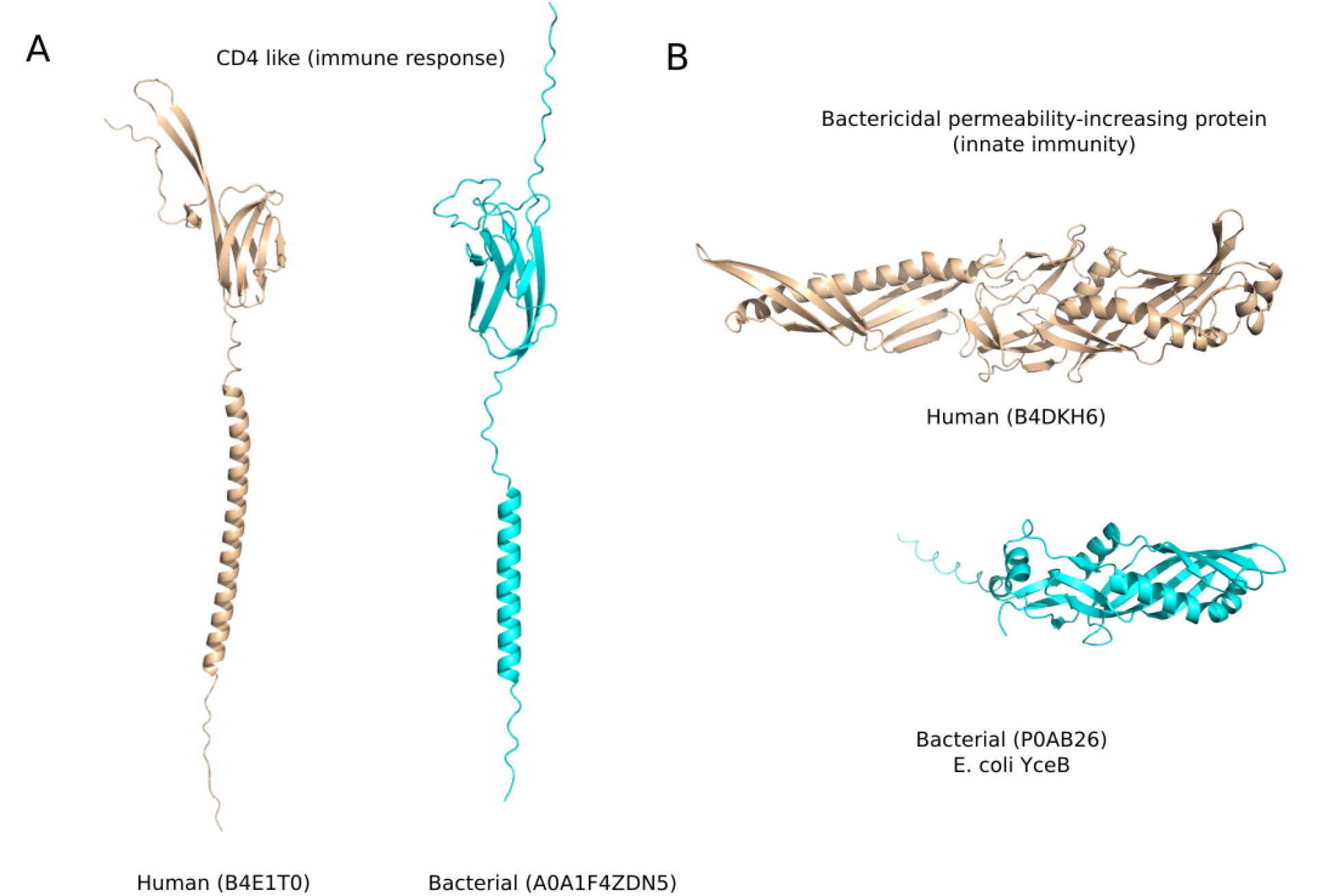
Additional examples of human related proteins in structural clusters with representatives or partial matches in bacterial species.

**Supplementary Figure 5.**
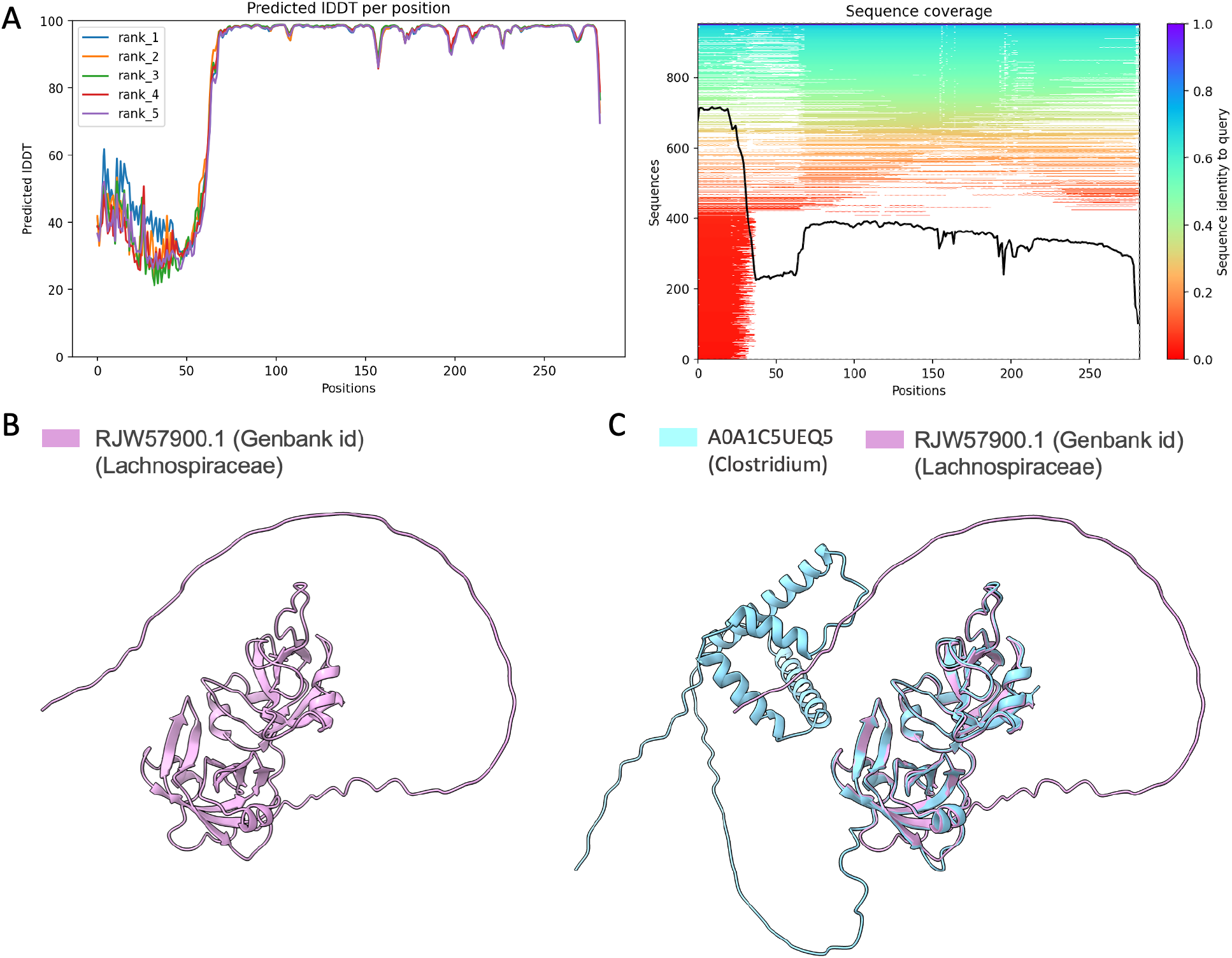
Comparison of predicted structures of homologous proteins: Lachnospiraceae bacterium to Clostridium. (A) pLDDT and multiple-sequence-alignment coverage output produced by ColabFold for the prediction of the protein sequence of *Lachnospiraceae*. (B) The predicted structure of RJW57900.1. (C) Superposition of the *Clostridium* protein structure with *Lachnospiraceae* with the DNA binding domain being well superposable.

